# EasyPseudogene: an easy-to-use and multithreaded pipeline for pseudogene detection

**DOI:** 10.64898/2026.03.04.709571

**Authors:** Chunhui Ai, Li Tan, Shenqiang Gao, Yong Wang

## Abstract

Pseudogenes are recognized as essential components for reconstructing adaptive evolutionary trajectories and understanding genomic remodeling. However, identifying these sequences in large eukaryotic genomes remains technically challenging due to fragmented workflows, complex manual configurations, and the lack of high-performance, parallelized tools capable of processing rapidly growing data volumes. We present EasyPseudogene, an automated and multithreaded pipeline designed for the end-to-end identification of pseudogenes across diverse eukaryotic lineages. Unlike traditional self-mapping tools that often fail to detect unitary pseudogenes when functional counterparts are absent, EasyPseudogene introduces an inter-species reference-driven paradigm that utilizes high-quality proteomes as probes to scan target genomes for evolutionary relics. The pipeline employs a modular “hierarchical screening and precision detection” architecture, integrating high-speed homology searching via MMseqs2 and spliced alignments via miniprot with high-fidelity, three-frame alignments using GeneWise. Performance benchmarking on cetacean genomes demonstrates that EasyPseudogene can replicate known gene loss events, such as the functional decay of the *ADRB3* gene, with 100% consistency relative to established manual workflows. By encapsulating complex comparative genomics logic into a standardized framework with interactive HTML visualization for mutation auditing at single-base resolution, EasyPseudogene provides a versatile and reproducible solution for marine ecology and evolutionary research.

## Introduction

In eukaryotic genomes, the dynamic interplay between gene gain and gene loss fundamentally shapes long-term adaptive processes in response to fluctuating environments (Coulombe-Huntington and Xia 2017; Deutekom et al. 2019). Gene loss and subsequent pseudogenization not only reflect the elimination of functional redundancy but also preserve an evolutionary record of historical selective pressures acting within specific ecological niches (Lopes-Marques et al. 2020; Chu et al. 2021). Pseudogenes are classically defined as genomic sequences that retain high sequence similarity to functional genes but have lost their protein-coding capacity due to disruptive mutations, including frameshifts and premature stop codons (Zheng and Gerstein 2007; Ranwez et al. 2011). With the rapid development of high-throughput sequencing technologies and large-scale comparative genomics, pseudogenes are no longer regarded as “junk DNA” but are increasingly recognized as informative components for elucidating genome evolutionary dynamics (Di Sanzo et al. 2020), gene regulatory network rewiring, and for improving the accuracy and completeness of genome annotation (An et al. 2017; Odah 2025).

Marine ecosystems impose extreme and heterogeneous selective pressures that frequently drive extensive genomic remodeling during biological adaptation (He et al. 2023; Szabó et al. 2026). Across diverse marine taxa—including mammals, fishes, and invertebrates—sensory systems, physiological regulatory pathways, and developmental processes often undergo systematic functional inactivation in response to lineage-specific ecological constraints (Yu et al. 2010; Wang et al. 2019; Zhang et al. 2021; Zhou et al. 2023). As a result, the comprehensive identification and characterization of pseudogenes in marine genomes has become essential not only for reconstructing adaptive evolutionary trajectories but also for establishing a robust foundation for cross-species comparative genomic analyses (Vigil-Stenman et al. 2015).

Existing software is characterized by pronounced taxonomic bias and functional fragmentation. Numerous tools are specialized for specific clades, such as pipelines optimized for human or prokaryote-specific utilities like PseudoPipe (Zhang et al. 2006) and Pseudofinder (Syberg-Olsen et al. 2022). Furthermore, many programs are restricted to specific pseudogene subtypes; for instance, PPFINDER and PΨFinder focus exclusively on processed pseudogenes derived from retrotransposition events (van Baren and Brent 2006; Abrahamsson et al. 2022). Consequently, there is a lack of a universal, open-source platform capable of transcending taxonomic boundaries to comprehensively address diverse evolutionary remnants.

The technical implementation of pseudogene identification is currently constrained by two major hurdles (Karro et al. 2007). First, existing workflows often rely on the manual configuration of heterogeneous software packages, characterized by complex installation procedures and fragmented dependencies, which lack a user-friendly, “one-click” automated pipeline for non-bioinformaticians. Second, when processing large eukaryotic genomes with high proportions of repetitive elements, many core algorithms operate via single-threaded or weakly parallelized mechanisms. This results in prohibitively long processing times—often spanning weeks—making it difficult to keep pace with the current trajectory of rapid data growth. Therefore, there is an urgent demand in the fields of comparative and ecological genomics for an integrated, easy-to-use, and fully multithreaded pseudogene identification pipeline.

In this study, high-quality cetacean genomes were selected as the benchmark datasets for performance evaluation. Cetaceans have undergone extensive gene inactivation events during their secondary transition from terrestrial to aquatic environments, making them an ideal model system for assessing the sensitivity and accuracy of pseudogene identification. It is important to emphasize that while cetaceans serve as a representative validation case, EasyPseudogene is designed for broad applicability across eukaryotic genomes, particularly marine non-model organisms characterized by complex genomic structures and limited baseline annotation. The pipeline is directly extensible to other marine mammals, fish, and invertebrates, providing a versatile and high-performance technical tool for marine ecology and evolutionary research.

## Methods

### Software architecture and implementation

EasyPseudogene was developed using a modular architecture designed to provide a fully automated pseudogene identification solution for large-scale eukaryotic genomes. The pipeline is driven by Python 3 and orchestrated via a series of Bash scripts (e.g., run_screening.sh, run_genewise.sh, and run_filtering.sh) to coordinate the execution of functional modules. To ensure reproducibility of the computational environment and streamline dependency management, all external core components are integrated into isolated environments managed via Conda (Grüning et al. 2018).

### Data collection and reference preparation

To evaluate the performance of the proposed pipeline, we utilized high-quality cetacean reference genomes, as benchmark datasets. Systematic identification within the pipeline is driven by three primary input components: 1. Reference Proteins (.fa): A curated set of protein sequences serves as the initial seeds for homology-based searching across the target assembly; 2. Target Genome (.fa): The complete whole-genome sequence designated for comprehensive functional annotation. 3. Configuration and Customization (Optional): To ensure flexibility and precision, the pipeline incorporates a default.conf file for the adjustment of stringent filtering thresholds. Additionally, users may provide a whitelist.txt to restrict the search space to specific gene families of interest.

### The Pre-GeneWise screening

To address the computational bottlenecks inherent in traditional whole-genome alignments, we implemented a hierarchical two-tier screening architecture for the rapid localization of candidate regions. Initially, MMseqs2 was employed for high-throughput, heuristic protein-to-genome searching (Fast Search) (Kallenborn et al. 2025). Subsequently, miniprot was utilized to perform spliced alignments, yielding precise exon-intron structural coordinates (Li 2023). Candidate sequences were prioritized based on alignment scores. To ensure the integrity of the genomic context, SAMtools was used to extract these regions, incorporating a default flanking length *L*_*flank*_ of 1000 bp upstream and downstream (Danecek et al. 2021). The genomic extraction interval *G*_*ext*_ was formally defined as:

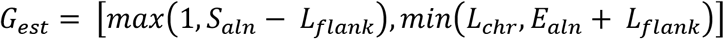

where *S*_*aln*_ and *E*_*aln*_ denote the start and end coordinates of the alignment, respectively, and *L*_*chr*_ represents the total length of the corresponding chromosome or scaffold.

### Detailed pseudogene detection and mutation analysis

Following the extraction of candidate genomic slices, the pipeline initiates the core phase of pseudogene detection and mutation characterization. To optimize computational throughput, the system implements a multi-threaded architecture that facilitates the parallel distribution of thousands of candidate regions across user-specified CPU resources. At the heart of this identification process, GeneWise is invoked to perform sophisticated protein-to-genome three-frame alignments (Birney et al. 2004). This approach enables the systematic recognition of disruptive mutational signatures—specifically premature stop codons and frameshift mutations—through exhaustive bidirectional searching on both the forward and reverse strands. Finally, all detected mutational features and alignment statistics are integrated and consolidated into a structured genewise.txt report, providing a comprehensive catalog of genomic features for downstream functional interpretation.

### Post-GeneWise filtering and quality evaluation protocols

To ensure maximum stringency and minimize the false discovery rate in the identification of pseudogenes, the EasyPseudogene pipeline implements a multi-stage filtering protocol encompassing six specialized sub-modules executed via the run_filtering.sh framework. The initial phase utilizes custom Python scripts to perform mutation counting and pre-filtering, thereby excluding candidate sequences that lack verifiable mutational signatures. To maintain the biological integrity of the dataset, unitary filters—comprising both Family and Redundancy filters—are applied to mitigate the confounding effects of multi-gene families and prevent interference from functional overlapping regions. Technical noise is further systematically mitigated through a multi-faceted false-positive exclusion process, which includes single-point mutation quality analysis, stringent homology thresholds, and end-mutation filtering designed to exclude stochastic errors occurring exclusively at sequence termini.

### Statistical analysis and performance metrics

The scalability of the pipeline was quantified by measuring total runtime and Peak Resident Set Size across varying CPU core counts during the processing of cetacean genomes. We performed a consistency analysis between the EasyPseudogene results and the original annotations published in Nature Communications, calculating Precision and Recall. For discordant loci, we further introduced outgroup species for synteny verification to confirm the authenticity of gene loss events.

## Results

### System architecture and engineering robustness compared to existing workflows

We implemented EasyPseudogene as an end-to-end automated pipeline designed to overcome the limitations of fragmented workflows and manual configurations. The system architecture is organized into five functional modules: input data management, hierarchical pre-screening, GeneWise-based precision analysis, multi-stage filtering, and standardized reporting (Fig. 1). Unlike traditional research that often relies on ad-hoc custom scripts with low reproducibility (Bai et al. 2019), EasyPseudogene provides a single-command execution framework that ensures seamless data orchestration and internal indexing (Table S1).

**Figure 1.**
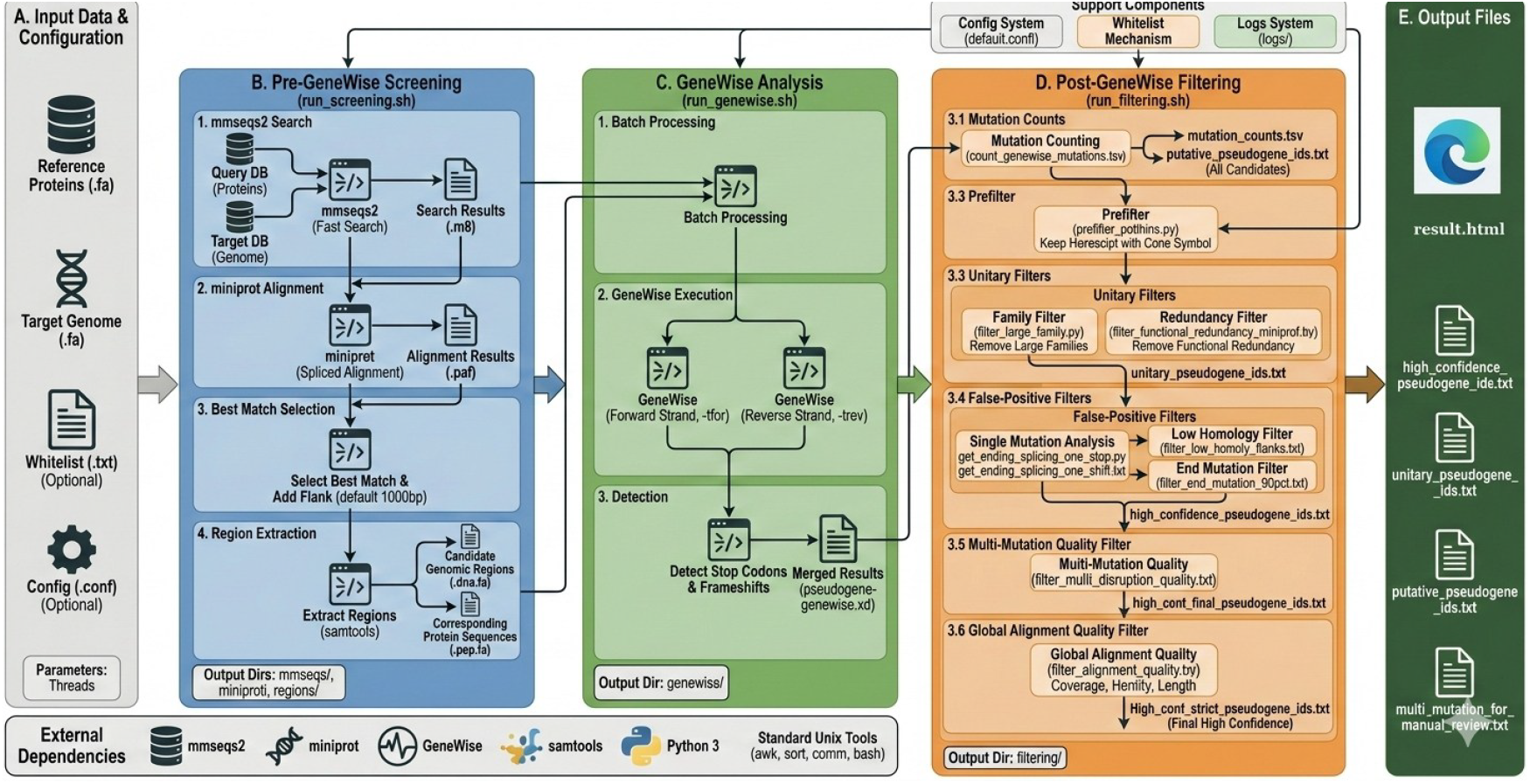
System architecture and implementation of the EasyPseudogene pipeline. The schematic illustrates the end-to-end automated workflow for pseudogene identification.

A comparative feature analysis demonstrates that EasyPseudogene is the only platform to integrate an inter-species reference-driven paradigm with HMM-based GeneWise alignment (Table S2). This architecture allows the pipeline to prioritize candidate regions through a two-tier screening process involving MMseqs2 and miniprot before performing exhaustive bidirectional searching for mutational signatures (Fig. 1). This engineering integration not only reduces the technical barrier for non-bioinformaticians but also establishes a rigorous foundation for cross-species pseudogene identification.

### Computational efficiency and scalability on large-scale genomic data

The performance of EasyPseudogene was quantified using genomic data at the scale of marine organisms to simulate real-world identification tasks. We measured execution time and memory usage across four orders of magnitude of input sequence counts (Fig. 2).

**Figure 2.**
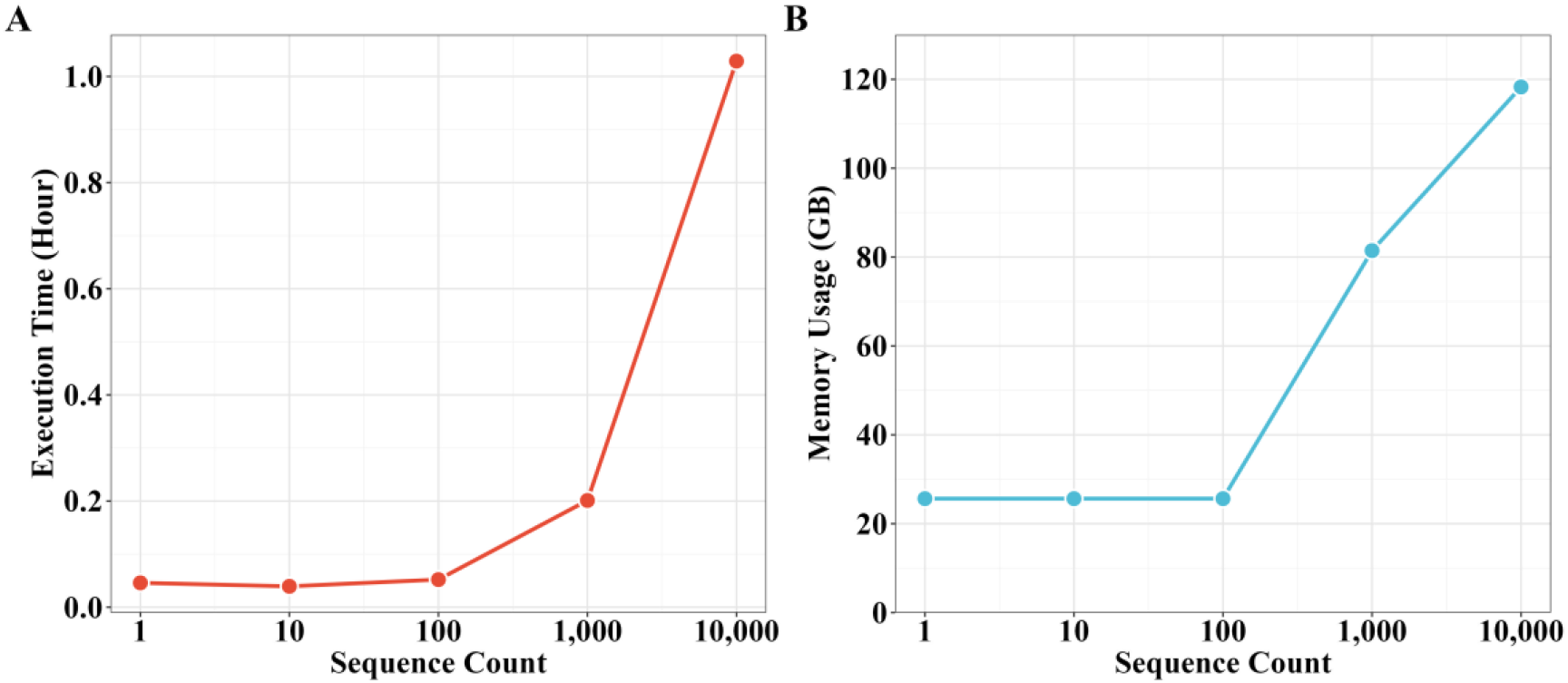
Computational performance and scalability of EasyPseudogene. (A) Execution time (hour) scaling as a function of the number of input protein sequences. (B) Peak memory usage (GB) recorded across varying sequence counts. Benchmarking was performed using mammalian-scale genomic data to simulate real-world large-scale identification tasks. The X-axis is presented on a log10 scale to visualize scaling across four orders of magnitude.

The pipeline exhibited high computational efficiency, with 1,000 sequences being processed in approximately 0.2 hours, and 10,000 sequences completed in 1.0 hour (Fig. 2A). Peak memory usage (Peak RSS) remained stable at approximately 25 GB for datasets of up to 100 sequences, scaling to 120 GB for the 10,000-sequence benchmark (Fig. 2B). These results demonstrate that EasyPseudogene can handle massive genomic datasets hundreds of times faster than manual orchestration, with the “time-to-first-view” determined primarily by CPU throughput.

### Sensitivity and biological validation through cetacean gene loss replication

To evaluate the recovery sensitivity of the pipeline, we utilized the longest human (Homo sapiens) protein sequences as query probes against the whole-genome assemblies of the sperm whale (Physeter macrocephalus) and dolphin (Tursiops truncatus). Using a benchmark of 90 curated high-quality transcripts, EasyPseudogene successfully identified 62 transcripts in the whale genome (68.9% sensitivity) and 65 in the dolphin genome (72.2% sensitivity). By integrating the detection results from both species, the workflow achieved a combined recovery rate of 88.9%, identifying 80 out of 90 benchmark transcripts (Fig. 3).

**Figure 3.**
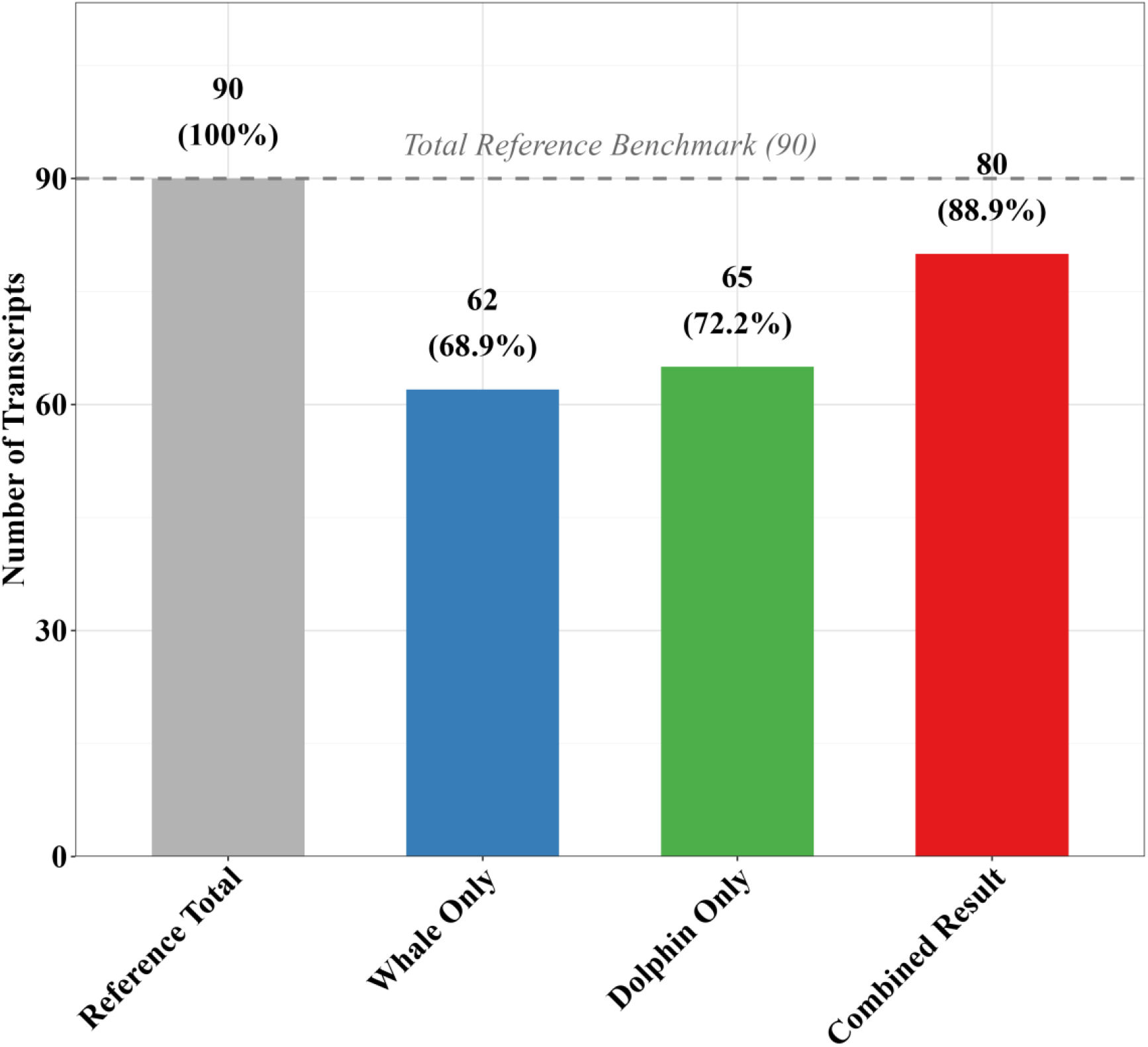
Performance validation and recovery sensitivity of EasyPseudogene. This bar chart illustrates the pipeline’s capability to recover high-quality pseudogene transcripts from the sperm whale (*Physeter macrocephalus*) and dolphin (*Tursiops truncatus*) genomes compared to a curated reference benchmark of 90 transcripts.

The precision of the automated detection was further validated by replicating the functional decay of the *ADRB3* gene (Transcript ID: ENST00000345060.5). By employing human protein transcripts as queries EasyPseudogene accurately localized a frameshift mutation at position 42 in the dolphin genome. This replication achieved 100% consistency with the results previously obtained through laborious manual workflows and custom scripts, demonstrating that the pipeline maintains rigorous analytical standards while significantly reducing technical barriers (Fig. 4).

**Figure 4.**
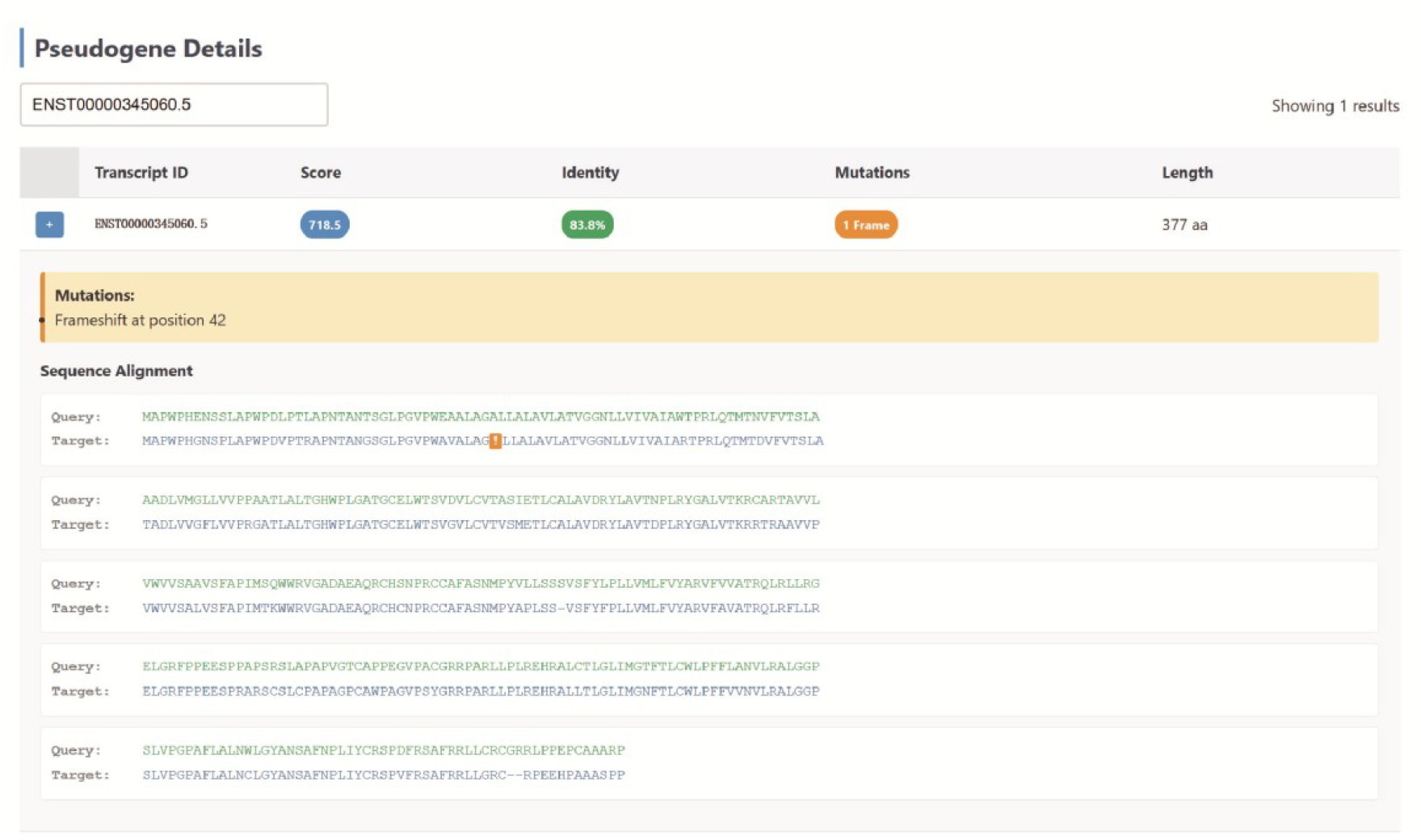
Interactive mutation auditing of the *ADRB3* gene at single-base resolution. Representative visualization of the functional decay in the *ADRB3* gene (Transcript ID: ENST00000345060.5).

### High-throughput landscape analysis and interactive dashboard

Beyond individual gene validation, EasyPseudogene provides a comprehensive landscape of pseudogenization events through its automated dashboard. In our performance trials, the system identified a total of 22,035 candidate regions, which were categorized into 417 regions with premature stop codons, 2,976 with frameshifts, and 93 harboring both mutation types (Fig. 5). The average alignment score across these identified relics was 838.0.

**Figure 5.**
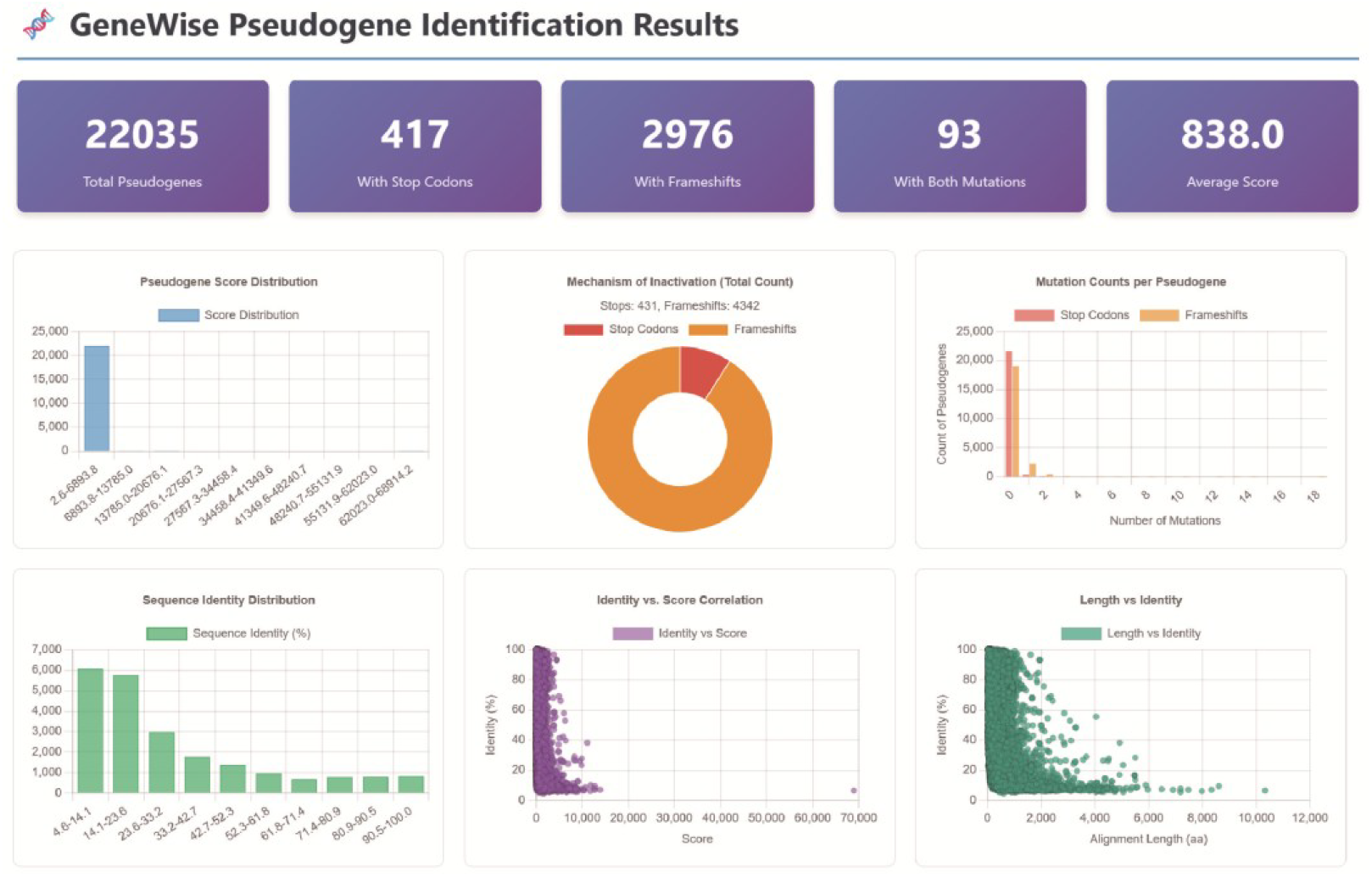
High-throughput pseudogene identification landscape and mutation profiles. The automated dashboard provides a comprehensive summary of the identified pseudogenes. Key visual metrics include the distribution of alignment scores and sequence identity, a quantitative breakdown of inactivation mechanisms, and correlation plots between alignment length and identity.

The interpretability of these large-scale datasets is enhanced by interactive HTML reports, allowing for the auditing of mutation sites at single-base resolution. This visualization enables the immediate verification of the molecular mechanisms underlying gene loss, transforming a traditionally time-consuming manual bottleneck into an intuitive, high-throughput verification process.

## Discussion

### A Paradigm Shift: From Intra-species Duplication to Inter-species Evolutionary Relics

The development of EasyPseudogene signifies a pivotal transformation in the methodology of pseudogene identification. Conventional identification tools, such as PseudoPipe, P-GRe, and PΨFinder, are primarily optimized for intra-species self-alignment, with their scientific objectives centered on identifying sequence copies generated via retrotransposition or gene duplication within a single genome. However, this paradigm faces inherent constraints when characterizing unitary pseudogenes (Zhang et al. 2010). When a gene undergoes complete functional inactivation and no longer yields a protein product in the target species, self-mapping approaches inevitably fail due to the absence of available “query seeds.”

In contrast, EasyPseudogene adopts an inter-species, reference-driven design paradigm. By leveraging high-quality reference proteomes as “probes,” the tool facilitates the precise localization of highly degraded functional relics across evolutionary timescales. This capability is paramount for reconstructing adaptive trajectories in response to extreme environmental shifts, such as the secondary aquatic transition of cetaceans (Hecker et al. 2017). The successful replication of the *ADRB3* gene loss event underscores the efficacy of this reference-driven strategy in elucidating the genomic basis of species-specific evolutionary traits. Furthermore, the integration of the GeneWise HMM model overcomes the alignment biases common to heuristic algorithms like BLAST when processing highly divergent sequences (Altschul et al. 1990; Eddy 1998), thereby ensuring single-nucleotide precision in capturing evolutionary breakpoints.

### Addressing the Reproducibility Crisis and Engineering Barriers in Evolutionary Genomics

A long-standing yet frequently overlooked challenge in contemporary evolutionary genomics is the “reproducibility crisis” (Dagli et al. 2024). Although numerous high-impact studies have identified critical gene loss events via ad-hoc custom scripts, the independent verification of these findings remains exceptionally difficult due to non-standardized workflows, fragmented data orchestration, and the general unavailability of source code. EasyPseudogene addresses this critical bottleneck by encapsulating complex comparative genomics logic into a standardized, end-to-end automated framework.

This engineering integration not only lowers the technical barrier for non-bioinformaticians but also guarantees analytical transparency through standardized configuration files. For instance, the implementation of a whitelist mechanism ensures that known evolutionary relics are preserved throughout large-scale filtering stages, while the interactive HTML reports transform previously laborious manual sequence auditing into an intuitive visual inspection process. In an era of escalating genomic data, the native support for high-thread-count parallelization enables researchers to conduct large-scale cross-species scans with minimal temporal cost. This transition from fragmented scripts to an engineered software solution not only enhances analytical efficiency but also establishes a rigorous foundation for constructing standardized, cross-species pseudogene databases in the future (Lam et al. 2009). With broad applicability to marine non-model organisms and other complex eukaryotes, EasyPseudogene is poised to become a versatile tool for resolving the genomic mechanisms underlying biological diversity.

## FUNDING

This work is supported by the National Natural Science Foundation of China (No. 42376149).

### Availability of data and materials

All the datasets supporting the conclusions have been presented within the article and its additional files. The source code for EasyPseudogene is publicly available on GitHub at https://github.com/chhhhai/EasyPseudogene.

## AUTHOR CONTRIBUTIONS

YW conceived and supervised the study and revised the manuscript. CHA developed the core pipeline and performed bioinformatics analysis. SQG collected and organized the datasets used for benchmarking. LT conducted the comprehensive software usability evaluation. The manuscript was prepared by CHA and YW.

## ETHICS STATEMENT

No animals or humans were involved in this study.

## CONSENT FOR PUBLICATION

All authors have approved the manuscript and given their consent for publication.

## References

Abrahamsson S, Eiengård F, Rohlin A, Dávila López M. 2022. PΨFinder: a practical tool for the identification and visualization of novel pseudogenes in DNA sequencing data. BMC Bioinformatics 23: 59.

Altschul SF, Gish W, Miller W, Myers EW, Lipman DJ. 1990. Basic local alignment search tool. Journal of Molecular Biology 215: 403–410.

An Y, Furber KL, Ji S. 2017. Pseudogenes regulate parental gene expression via ceRNA network. Journal of Cellular and Molecular Medicine 21: 185–192.

Bai L, Liu B, Ji C, Zhao S, Liu S, Wang R, Wang W, Yao P, Li X, Fu X et al. 2019. Hypoxic and Cold Adaptation Insights from the Himalayan Marmot Genome. iScience 11: 519–530.

Birney E, Clamp M, Durbin R. 2004. GeneWise and genomewise. Genome Res 14: 988–995.

Chu X, Li S, Wang S, Luo D, Luo H. 2021. Gene loss through pseudogenization contributes to the ecological diversification of a generalist Roseobacter lineage. Isme j 15: 489–502.

Coulombe-Huntington J, Xia Y. 2017. Network Centrality Analysis in Fungi Reveals Complex Regulation of Lost and Gained Genes. PLoS One 12: e0169459.

Dagli N, Haque M, Kumar S. 2024. The Replication Crisis: A Persistent Challenge in Biomedical Research. Bangladesh Journal of Medical Science 23: 907–910.

Danecek P, Bonfield JK, Liddle J, Marshall J, Ohan V, Pollard MO, Whitwham A, Keane T, McCarthy SA, Davies RM et al. 2021. Twelve years of SAMtools and BCFtools. GigaScience 10.

Deutekom ES, Vosseberg J, van Dam TJP, Snel B. 2019. Measuring the impact of gene prediction on gene loss estimates in Eukaryotes by quantifying falsely inferred absences. Plos Computational Biology 15.

Di Sanzo M, Quaresima B, Biamonte F, Palmieri C, Faniello MC. 2020. FTH1 Pseudogenes in Cancer and Cell Metabolism. Cells 9.

Eddy SR. 1998. Profile hidden Markov models. Bioinformatics 14: 755–763.

Grüning B, Dale R, Sjödin A, Chapman BA, Rowe J, Tomkins-Tinch CH, Valieris R, Köster J. 2018. Bioconda: sustainable and comprehensive software distribution for the life sciences. Nat Methods 15: 475–476.

He X, Wang H, Xu T, Zhang Y, Chen C, Sun Y, Qiu JW, Zhou Y, Sun J. 2023. Genomic Analysis of a Scale Worm Provides Insights into Its Adaptation to Deep-Sea Hydrothermal Vents. Genome Biol Evol 15.

Hecker N, Sharma V, Hiller M. 2017. Transition to an Aquatic Habitat Permitted the Repeated Loss of the Pleiotropic KLK8 Gene in Mammals. Genome Biol Evol 9: 3179–3188.

Kallenborn F, Chacon A, Hundt C, Sirelkhatim H, Didi K, Cha S, Dallago C, Mirdita M, Schmidt B, Steinegger M. 2025. GPU-accelerated homology search with MMseqs2. Nature Methods 22: 2024–2027.

Karro JE, Yan Y, Zheng D, Zhang Z, Carriero N, Cayting P, Harrrison P, Gerstein M. 2007. Pseudogene.org: a comprehensive database and comparison platform for pseudogene annotation. Nucleic Acids Res 35: D55–60.

Lam HY, Khurana E, Fang G, Cayting P, Carriero N, Cheung KH, Gerstein MB. 2009. Pseudofam: the pseudogene families database. Nucleic Acids Res 37: D738–743.

Li H. 2023. Protein-to-genome alignment with miniprot. Bioinformatics 39.

Lopes-Marques M, Serrano C, Cardoso AR, Salazar R, Seixas S, Amorim A, Azevedo L, Prata MJ. 2020. GBA3: a polymorphic pseudogene in humans that experienced repeated gene loss during mammalian evolution. Scientific Reports 10: 11565.

Odah MAA. 2025. The Dark Genome Investigating Pseudogenes and Non-Coding Regions in Genetic Regulation. Preprints doi:10.20944/preprints202506.2471.v1.

Ranwez V, Harispe S, Delsuc F, Douzery EJ. 2011. MACSE: Multiple Alignment of Coding SEquences accounting for frameshifts and stop codons. PLoS One 6: e22594.

Syberg-Olsen MJ, Garber AI, Keeling PJ, McCutcheon JP, Husnik F. 2022. Pseudofinder: Detection of Pseudogenes in Prokaryotic Genomes. Mol Biol Evol 39.

Szabó G, Eloe-Fadrosh EA, Pett-Ridge J, Woyke T. 2026. A genomic view of Earth’s biomes. Nature Reviews Genetics 27: 13–31.

van Baren MJ, Brent MR. 2006. Iterative gene prediction and pseudogene removal improves genome annotation. Genome Res 16: 678–685.

Vigil-Stenman T, Larsson J, Nylander JA, Bergman B. 2015. Local hopping mobile DNA implicated in pseudogene formation and reductive evolution in an obligate cyanobacteria-plant symbiosis. BMC Genomics 16: 193.

Wang K, Shen Y, Yang Y, Gan X, Liu G, Hu K, Li Y, Gao Z, Zhu L, Yan G et al. 2019. Morphology and genome of a snailfish from the Mariana Trench provide insights into deep-sea adaptation. Nature Ecology & Evolution 3: 823–833.

Yu L, Jin W, Wang J-x, Zhang X, Chen M-m, Zhu Z-h, Lee H, Lee M, Zhang Y-p. 2010. Characterization of TRPC2, an Essential Genetic Component of VNS Chemoreception, Provides Insights into the Evolution of Pheromonal Olfaction in Secondary-Adapted Marine Mammals. Mol Biol Evol 27: 1467–1477.

Zhang Y, Wang J, Lv M, Gao H, Meng L, Yunga A, Seim I, Zhang H, Liu S, Zhang L et al. 2021. Diversity, function and evolution of marine invertebrate genomes. bioRxiv doi:10.1101/2021.10.31.465852: 2021.2010.2031.465852.

Zhang Z, Carriero N, Zheng D, Karro J, Harrison PM, Gerstein M. 2006. PseudoPipe: an automated pseudogene identification pipeline. Bioinformatics 22: 1437–1439.

Zhang ZD, Frankish A, Hunt T, Harrow J, Gerstein M. 2010. Identification and analysis of unitary pseudogenes: historic and contemporary gene losses in humans and other primates. Genome Biology 11.

Zheng D, Gerstein MB. 2007. The ambiguous boundary between genes and pseudogenes: the dead rise up, or do they? Trends in Genetics 23: 219–224.

Zhou Y, Liu H, Feng C, Lu Z, Liu J, Huang Y, Tang H, Xu Z, Pu Y, Zhang H. 2023. Genetic adaptations of sea anemone to hydrothermal environment. Sci Adv 9: eadh0474.

